# A virtual B cell lymphoma model to predict effective combination therapy

**DOI:** 10.1101/037093

**Authors:** Wei Du, Rebecca Goldstein, Yanwen Jiang, Omar Aly, Leandro Cerchietti, Ari Melnick, Olivier Elemento

## Abstract

**Abstract:** The complexity of cancer signaling networks limits the efficacy of most single agent treatments and brings about challenges in identifying effective combinatorial therapies. Using chronic active B cell receptor (BCR) signaling in diffuse large B cell lymphoma (DLBCL) as a model system, we established a computational framework to optimize combinatorial therapy *in silico.* We constructed a detailed kinetic model of the BCR signaling network, which captured the known complex crosstalk between the NFκB, ERK and AKT pathways; and multiple feedback loops. Combining this signaling model with a data-derived tumor growth model we predicted viability responses of many single drug and drug combinations that are in agreement with experimental data. Under this framework, we exhaustively predicted and ranked the efficacy and synergism of all possible combinatorial inhibitions of eleven currently targetable kinases in the BCR signaling network. Our work established a detailed kinetic model of the core BCR signaling network and provides the means to explore the large space of possible drug combinations.

**Author Summary:** Using chronic active B cell receptor (BCR) signaling in diffuse large B cell lymphoma(DLBCL) as a model system, we developed a kinetic-modeling based computational framework for predicting effective combination therapy *in silico.* By integrative modeling of signal transduction, drug kinetics and tumor growth, we were able to directly predict drug-induced cell viability response at various dosages, which were in agreement with published cell line experimental data. We implemented computational screening methods that identified potent and synergistic combinations *in silico* and validated our independent predictions *in vitro.*

## Introduction

The activation of intracellular signaling pathways in response to environmental stimulus leads to important cell decisions such as proliferation. The amplitude and duration of pathway activation are precisely and robustly controlled by complex regulatory loops to maintain cellular homeostasis. In cancer, activating mutations or deletion of signaling repressors frequently result in sustained and exaggerated pathway activation that drives uncontrolled tumor survival and proliferation. Targeted therapies that use small molecule inhibitors to repress specific signaling pathway members, e.g. kinases, can directly block oncogenic pathway activation and lead to tumor cell death. These targeted therapies are expected to provide improved efficacy and reduced toxicity compared to chemotherapy.

However, clinical application of targeted therapy is facing several challenges such as low response rate and frequently acquired drug resistance. The limited efficacy of single agent targeted therapy is at least partially due to pathway crosstalks and compensatory circuits within signaling networks targeted by these agents[1]. Crosstalks and compensatory circuits allow signals to bypass drug inhibition and reactivate downstream effectors. By simultaneously repressing multiple nodes in a signaling network, combination therapy has the potential to completely extinguish oncogenic signaling and induce more potent and durable treatment response. Thus, novel drug combinations where two or more drugs work cooperatively to suppress corrupted signaling networks need to be identified to achieve maximum therapeutic efficacy. The complexity of signaling networks makes it difficult to simply guess which combinations will be effective and synergistic and which ones will not. Moreover, given the large number of possible drug combinations against complex signaling networks, comprehensive experimental screening - including exploration of multiple dosages - is not practically feasible.

Besides, results acquired from such screening may be specific to the cell line tested, thus lacking general applicability to highly variable primary tumors found in patients.

Computational models of signaling networks that can accurately reconstruct signaling dynamics *in silico* may represent a useful alternative to experimental screening and trial-and-error experimental investigation. Once proven reliable, these models can be used to exhaustively test the efficacy of a large number of single drug and drug combinations by quantifying signaling output under corresponding network perturbations. Even though computational modeling has been widely used to study the dynamics of signaling network in the past decades, the development of cancer signaling models and its application to predicting effective combinatorial therapies is still lacking. Here we demonstrate the feasibility of this approach using chronic activation of B cell receptor (BCR) signaling in diffuse large B cell lymphoma (DLBCL) as a model system. We adopted a systems biology approach and established a computational framework to optimize anti-DLBCL combinatorial therapy *in silico.* The proposed approach is broadly applicable and can be used for other malignancies driven by aberrantly active signaling pathways.

The deregulation of B cell receptor (BCR) signaling is central to the pathogenesis of many B cell malignancies. It is especially central in the activated B cell-like subtype of diffuse large B cell lymphoma (ABC-DLBCL). ABC DLBCLs exhibit chronic active BCR signaling, and are addicted to constitutive activation of downstream survival and proliferation signals such as NFκB[2]. It has recently been found that a subset of the germinal center B cell like(GCB) subtype of DLBCLs are also dependent on BCR signaling through activation of the PI3K/AKT pathway [3]. Multiple small molecule inhibitors against BCR signaling were developed and proved effective in killing BCR-dependent DLBCLs *in vitro* and *in vivo* [4,5,6]. However when tested in clinical trials, single agent treatments again demonstrated limited responsiveness and efficacy[7], suggesting an urgent need for the design of effective combination therapies.

In this work, we present a kinetic modeling-based computational framework for predicting and optimizing combinatorial therapy against chronic active BCR signaling (Fig 1). We constructed a detailed kinetic model of the BCR signaling network parameterized by published signaling responses and protein concentrations. Mathematical models of proximal BCR signaling and downstream transcriptional network have been reported [8,9,10]. But to our knowledge, this is the first kinetic model to reconstruct the entire core BCR signaling network *in silico.* Using published drug response data in a BCR signaling-dependent cell line, we trained a tumor growth model which in combination with the kinetic model allowed us to simulate viability response upon various targeted treatments. Under this framework, we exhaustively tested the efficacy and synergism of all possible combinations of inhibition of eleven currently targetable kinases in the BCR signaling network. We discuss how these results pave the way for the discovery of effective drug combinations.

**Fig 1.**
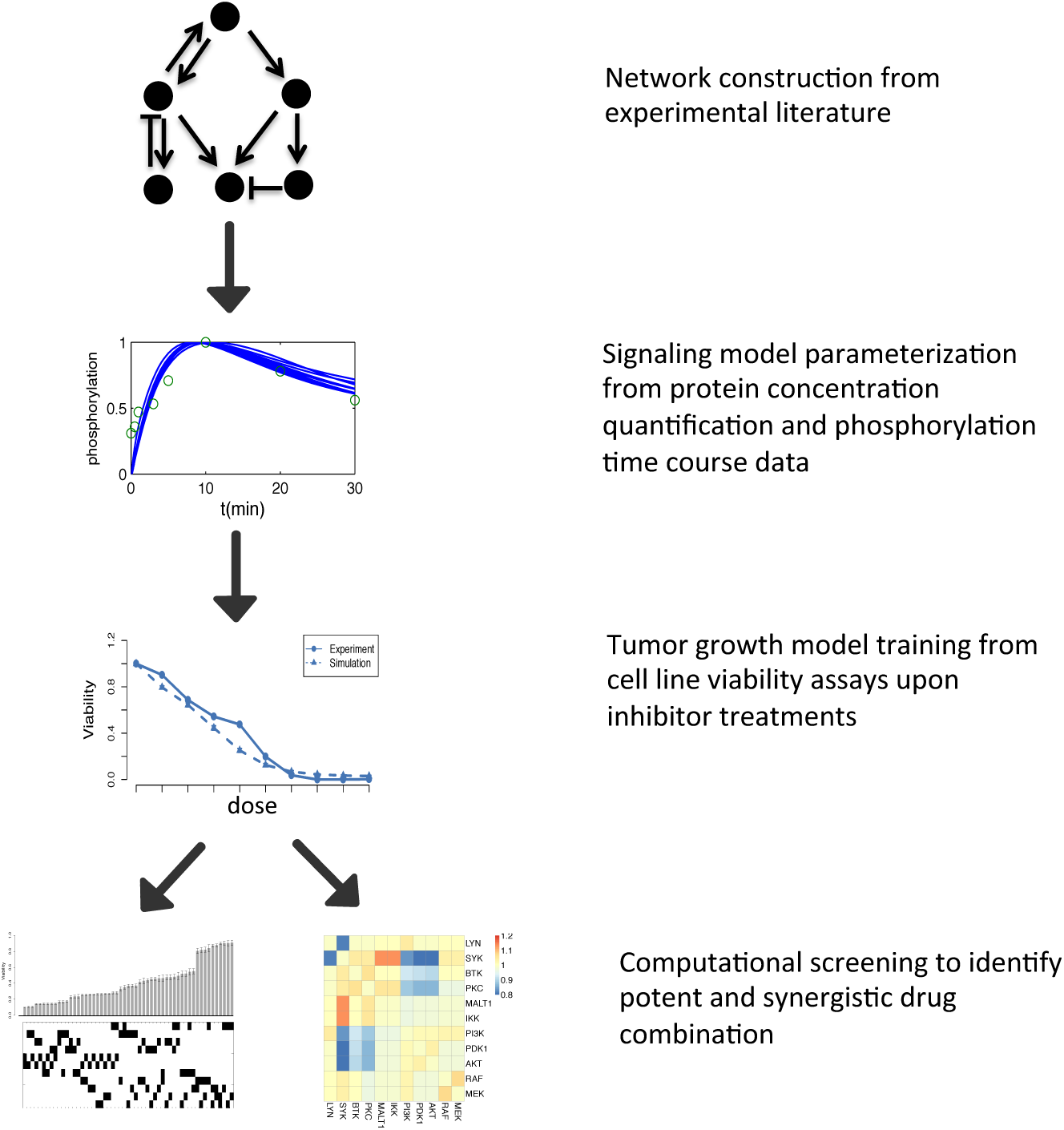
Outline of the approach taken in the present study. The central BCR signaling network was constructed based on validated protein-protein interactions from experimental literature. Parameters of molecular reaction kinetics were estimated from phosphorylation time course data and protein concentrations were retrieved from MOPED protein expression database. A phenotypic tumor growth model was trained on cell viability assays of inhibitor treatments to link signaling response to viability outcome. In the end, simulation of the signaling model in combination with the tumor growth model was conducted to optimize treatment strategy. The model’s prediction was compared to published drug response data and new prediction-driven hypotheses were tested independently in vitro.

## Results

### Kinetic modeling of BCR signaling network reproduces normal BCR signaling *in silico*

We first curated the central BCR signaling network by gathering experimentally validated protein-protein interactions from literature. The reconstructed network is shown in Fig 2, and includes three major signaling pathways downstream of BCR, namely NFκB, PI3K/AKT and RAF/RAS/ERK. We chose to include these three pathways because they have been reported to closely regulate cell survival and proliferation in B cells and B cell malignancies [11]. Antigen-induced BCR crosslinking allows SRC family kinases, mainly LYN, to phosphorylate the immuno-receptor tyrosine based activation motifs (ITAMs) of the intracellular BCR subunits Igα (CD79A) and Igβ (CD79B)[12]. Dually phosphorylated ITAM motifs then recruit SYK and activate it via tyrosine phosphorylation[13]. Activated SYK phosphorylates adapter BLNK, which recruits BTK to the plasma membrane to facilitate its phosphorylation and subsequent activation by SYK and LYN[14]. Activated BTK further phosphorylates PLCγ2, which catalyzes the hydrolysis of phosphatidylinositol-4,5-bisphosphate (PI(4,5)P_2_) into diacylglycerol (DAG) and inositol trisphosphate (IP_3_)[15]. DAG together with elevated intracellular calcium induced by IP_3_ triggered endoplasmic reticulum (ER) calcium release activates PKCβ[16], which then stimulates two diverge pathways that activate NFκB and ERK respectively. Phosphorylation of CARD11 by PKCβ leads to the assembly of the CBM complex composed of CARD11, BCL10 and MALT1[17]. CBM acts as a scaffolding complex that facilitates IKK phosphorylation by TAK1, which in turn phosphorylates IKB and induces its degradation, releasing NFκB into the nucleus to elicit transcriptional activity[18]. Additionally, protease activity of MALT1 positively regulates NFκB signaling by cleaving and inactivating inhibitors against NFκB activation such as A20 and RELB[19,20]. In the meantime, PKCβ and DAG activate RASGRP, which triggers the canonical MAPK signaling cascade, leading to eventual phosphorylation and activation of ERK[21]. On the other hand, SYK and LYN phosphorylate BCAP and CD19 respectively, which activate PI3K by membrane recruitment[22,23]. PI(3,4,5)P_3_ synthesized by PI3K further facilitates PDK1 catalyzed AKT phosphorylation by binding to both proteins via their plextrin homology (PH) domains[24]. Importantly, LYN negatively regulates PI3K signaling by activating SHIP1, which hydrolyzes PI(3,4,5)P_3_ into PI(4,5)P_2_[25].

**Fig 2.**
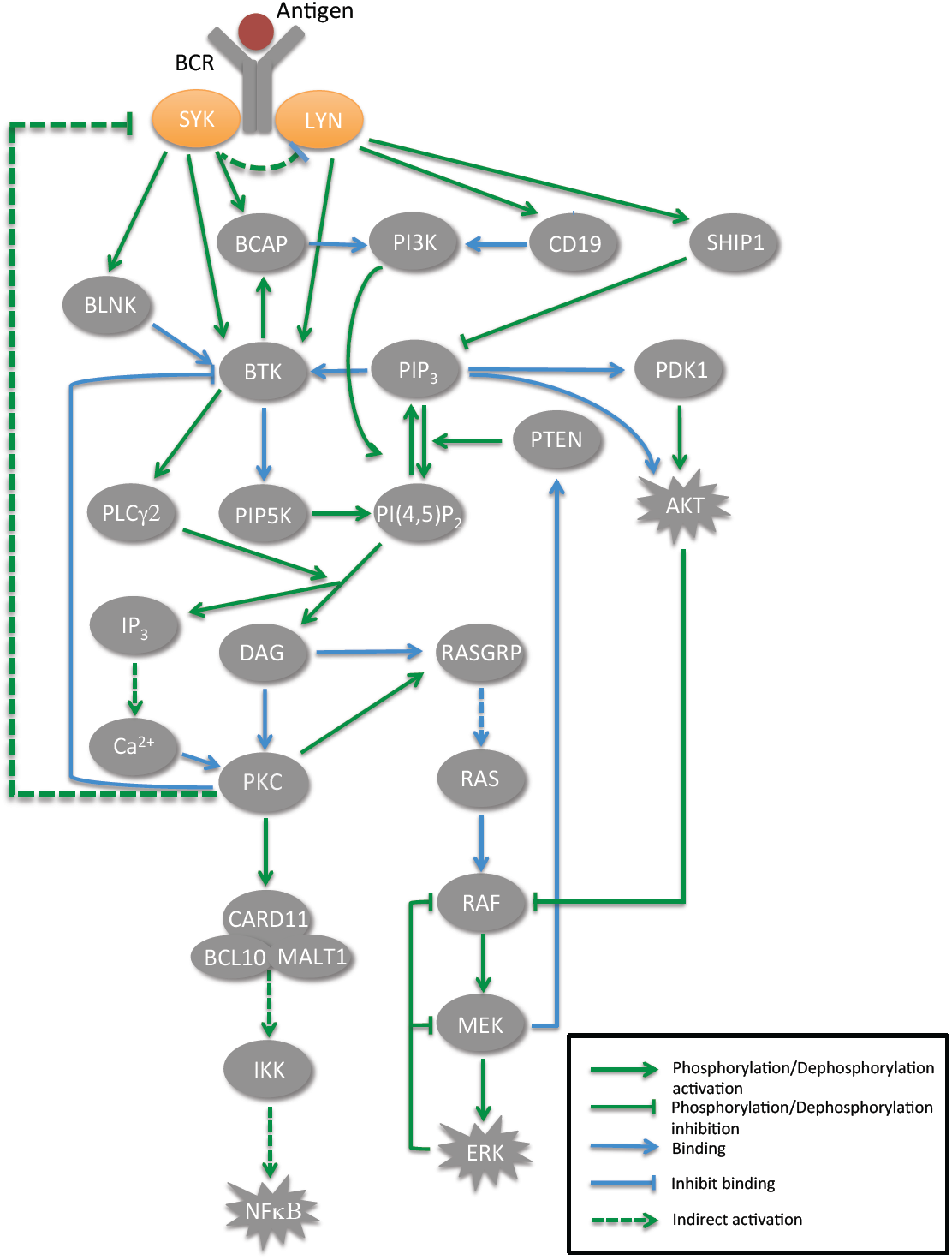
The central BCR signaling network constructed from literature. Antigen binding induces BCR aggregation and subsequent phosphorylation, which further triggers a complex signaling cascade initiated by phosphorylated LYN and SYK. The BTK-PLCγ2-PKCβ pathway activates downstream NFκB and ERK through divergent paths, while membrane recruitment of PI3K leads to AKT activation. Pathway crosstalks and feedback regulations are highly abundant in the network.

Besides major signal transduction pathways as described above, our model includes key regulatory interactions in the BCR signaling network such as pathway crosstalks and feedback loops. The PI3K pathway positively regulates NFκB and ERK signaling by enhancing BTK membrane recruitment via PI(3,4,5)P_3_ binding. In the meantime, it conversely attenuates ERK signaling via AKT catalyzed RAF phosphorylation[26]. It has recently been found that MEK negatively regulates PI3K/AKT signaling by recruiting PTEN to the plasma membrane [27], which dephosphorylates PI(3,4,5)P_3_ into PI(4,5)P_2_. BTK amplifies BCR signaling by two coupled positive feedback loops. It recruits PIP5K to the plasma membrane, which produces PI(4,5)P_2_ to sustain both PI(3,4,5)P_3_ synthesis and PI(4,5)P_2_ hydrolysis[28]. Additionally, BTK phosphorylates BCAP, further facilitating the membrane recruitment of PI3K[23]. The activity of BTK is attenuated by active PKCβ via disruption of its membrane localization, constituting a negative feedback loop[29]. Besides, another indirect feedback from PKC to SYK was added into the model as knockdown of PKCδ was shown to mediate hyperphosphorylation of SYK[30]. Furthermore, multiple negative feedback loops exist within the MAPK signaling cascade to fine-tune its activation amplitude and duration[31].

Instead of directly applying mass action kinetics to characterize elementary reactions in the network, we chose to adopt more streamlined mathematical representations derived from mass action law under reasonable assumptions **(see Materials and Methods)**. This strategy greatly reduced the number of variables, equations and most importantly parameters required in the mathematical model. As elementary protein-protein binding reactions generally reach equilibrium within seconds, we modeled them by deriving the steady-state relationships from mass action law **(see Materials and Methods)**. For enzymatic reactions such as phosphorylation or dephosphorylation, we adopted a classic Michaelis-Menten kinetic framework. To parameterize the model, we first retrieved protein concentrations in B lymphocytes quantified by mass-spectrometry from the MOPED protein expression database [32]. We modeled LYN and SYK as two independent input signals that triggered a downstream response. Their activation kinetics were approximated by double exponential functions (see **Materials and Methods**) where parameters in the functions were estimated by fitting to the phosphorylation time course data[33]. We then used genetic algorithms to optimize the remaining 72 kinetic parameters within bounded biologically reasonable ranges (**S1 Table**) by minimizing residual sum of squares between simulated phosphorylation time courses and published western blot data[33]. Experimental data and simulated results were each normalized to their respective maximum value for comparison. 10 sets of kinetic parameters were identified from 5000 independent runs that fit almost equally well. Simulated trajectory under these 10 parameter sets together with phosphorylation time course data are shown in Fig 3A. Parameter sensitivity analysis was performed as described in **Materials and Methods (S1 Fig)**. Of note, we were independently able to find 39 kinetic parameter values from the literature, and we compared these values with the range of estimated 10 parameter sets (Fig 3B). By shuffling the literature-retrieved parameters 10,000 times, we found that the literature-retrieved parameter values fall within the estimated parameter ranges significantly more often than random (p=0.05)(Fig 3C, 3D). We note however that many discrepancies were found between estimated and published parameters (Fig 3B). We speculate that many of these discrepancies are likely due to *in vitro* nature of the experiments used to quantify kinetic parameters.

**Fig 3.**
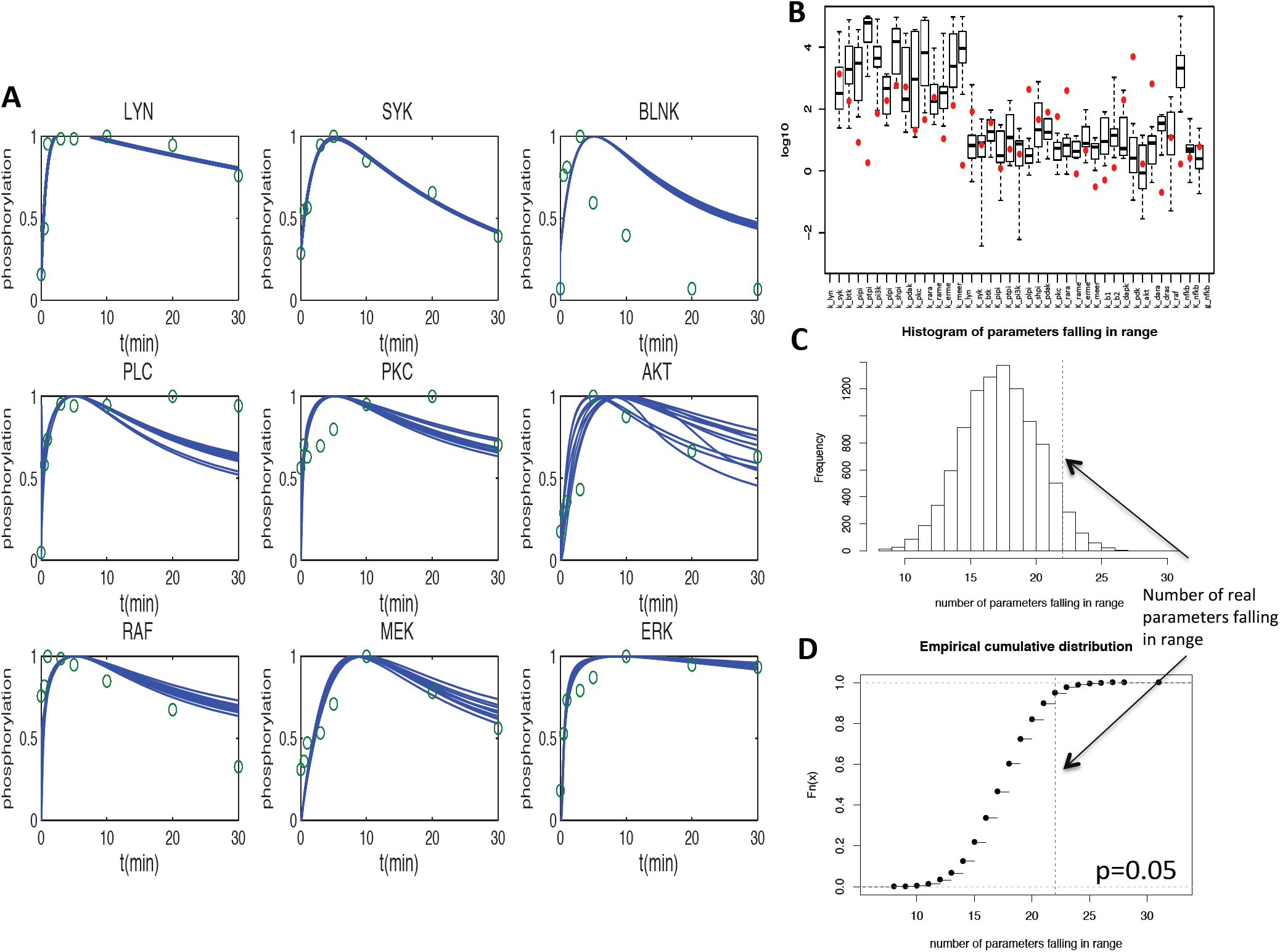
Simulation of normal BCR signaling and estimation of kinetic parameters. (**A**) Simulated trajectory of ten parameter sets in comparison with published phosphorylation time course data. (**B**) Comparison between literature-retrieved parameter values with simulation-estimated parameter ranges. Box-plot indicates the simulation-estimated parameter ranges while red dots represent literature-retrieved parameter values. (**C**) Shuffling of the literature-retrieved parameters 10,000 times to obtain a distribution of number of parameters that would fall within simulation-estimated parameter ranges by chance. (**D**) Empirical cumulative distribution of the number of parameters that would fall within simulation-estimated parameter ranges by chance.

### Combining BCR signaling model with a tumor growth model predicted cell viability response upon single and combinatorial drug treatments in a BCR signaling-dependent ABC-DLBCL cell line

We next sought to simulate the effect of various small molecule inhibitors on ABC DLBCL cell viability and to compare simulation results with published drug response data in a BCR signaling dependent ABC-DLBCL cell line TMD8 [34]. We selected TMD8 because of the extensive drug combinatorial data available on this cell line[34]. We first made several modifications to the model to accommodate the differences between normal BCR signaling and aberrant BCR signaling in ABC-DLBCL. Instead of applying a temporal upstream stimulus, we assumed constitutive LYN and SYK phosphorylation as observed both in ABC DLBCL cell lines and in primary DLBCL patient samples [2,35] **(see Materials and Methods)**. Additionally, we accounted for genetic alterations in members of the BCR signaling network in TMD8 compared to normal B cells. Specifically, TMD8 was shown to carry CD79B mutation that attenuates LYN activity by approximately 80%[2]. Correspondingly we decreased the enzymatic activity of LYN in the model to the same extent **(see Materials and Methods)**.

To predict cell viability response from signaling output, we formulated a tumor growth model in which the growth rate of tumor cells is dependent on the weighted sum of the three downstream survival and proliferation signals NFκB, ERK and AKT through a Hill function **(see Materials and Methods)**. Similar formalism has been used to characterize tumor growth of ERBB-amplified breast cancer driven by ERK and AKT activation [36]. We used published viability response data in TMD8 to parameterize the tumor growth model, where cells were treated with IKK, AKT and MEK inhibitors at multiple dosages[34]. Specifically, using the median effect equation [37], we calibrated the percent activity left on the targeted kinase for each inhibitor at a given dosage based on the inhibitor’s IC_50_ value **(see Materials and Methods, S2 Table)**. We then reduced the activity of the targeted kinase to the same level in the model and simulated steady-state signaling output. Parameters in the tumor growth model were estimated by minimizing residual sum of squares between predicted viability response and experimental data.

We first simulated single drug viability response of inhibitors covering the NFκB, PI3K/AKT and MAPK pathway and compared to experimental data. We observed that *in silico* simulation with the BCR signaling model and the tumor growth model recapitulated the viability response of the three training single drug response, namely IKK, AKT, MEK inhibitors [34] (Fig 4A). This is not surprising since the growth model was fitted based on training data. As independent predictions, we also simulated drug response of inhibitors targeting other kinases in the network, e.g. CAL-101 against PI3K, Ibrutinib against BTK, and found that predicted viability response matched favorably with TMD8 drug response data[34] (Fig 4B). At the same time, we found simulated viability response of SYK inhibition to deviate from experimental data(grey line), yet this discrepancy can be partially rescued by adding a negative feedback from SYK to LYN(blue line). It has been reported that SYK functions as a negative regulator of BCR signaling by phosphorylating Ig-α[30,38,39]. Since Ig-α primarily interacts with LYN, we assumed in the model that SYK indirectly negatively regulates LYN.

**Fig 4.**
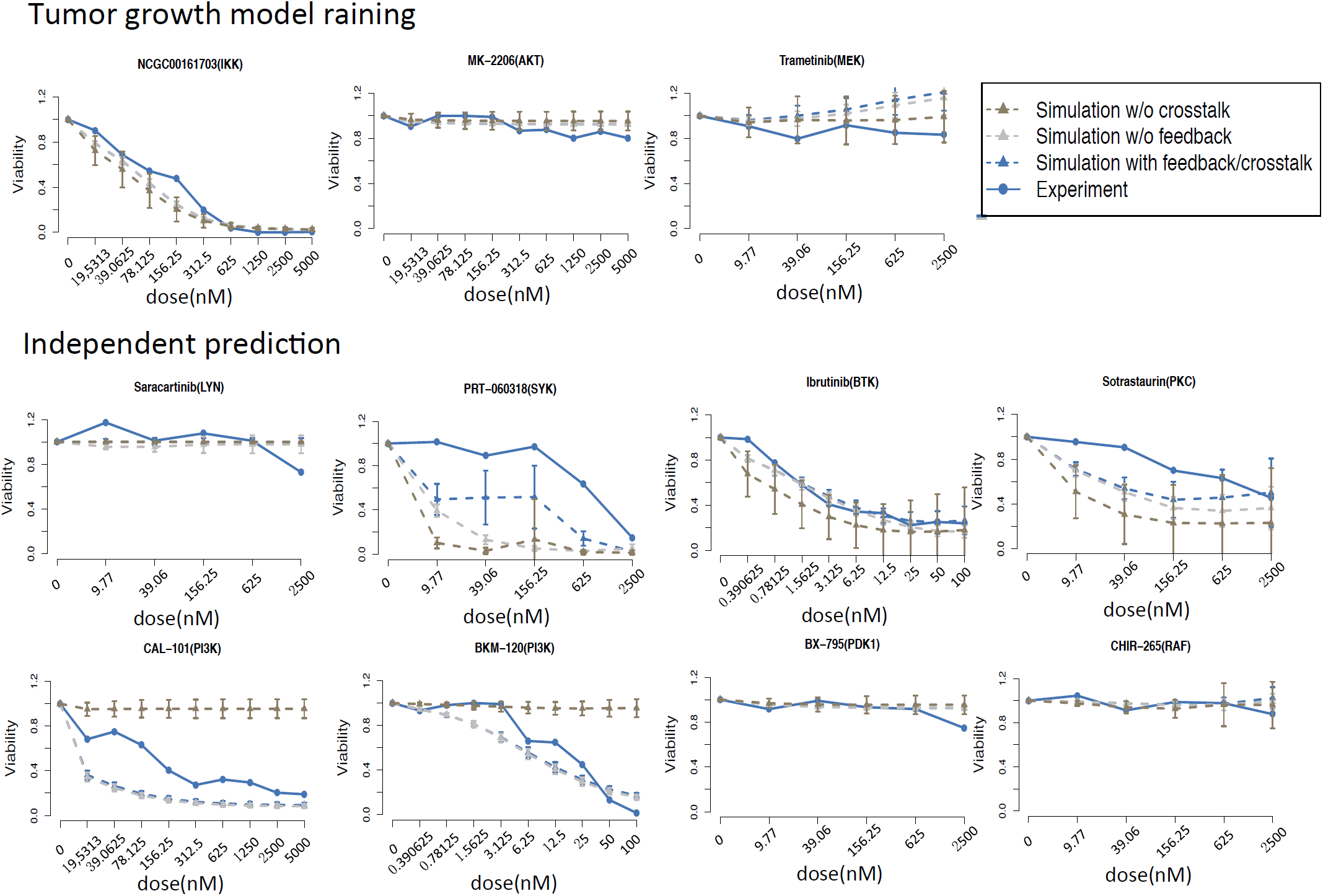
Training and prediction of single drug viability response in ABC DLBCL cell line TMD8. (**A**) Tumor growth model parameterization using single drug viability response of inhibitors targeting NFκB, AKT and MEK. Gray dashed lines correspond to simulation results of model without SYK to LYN negative feedback, while brown dashed lines correspond to simulation results of model without PI3K-NFκB crosstalk. (**B**) Single drug viability response of inhibitor targeting various kinases against BCR signaling network.

Beyond single drug viability response, we also simulated combinatorial drug response of Ibrutinib in combination with various other kinases targeting the BCR pathway, and observed the predicted response contour to match favorably with experimental results(Fig 5). These results demonstrate that our model can correctly capture the interaction between inhibitors as well.

**Fig 5.**
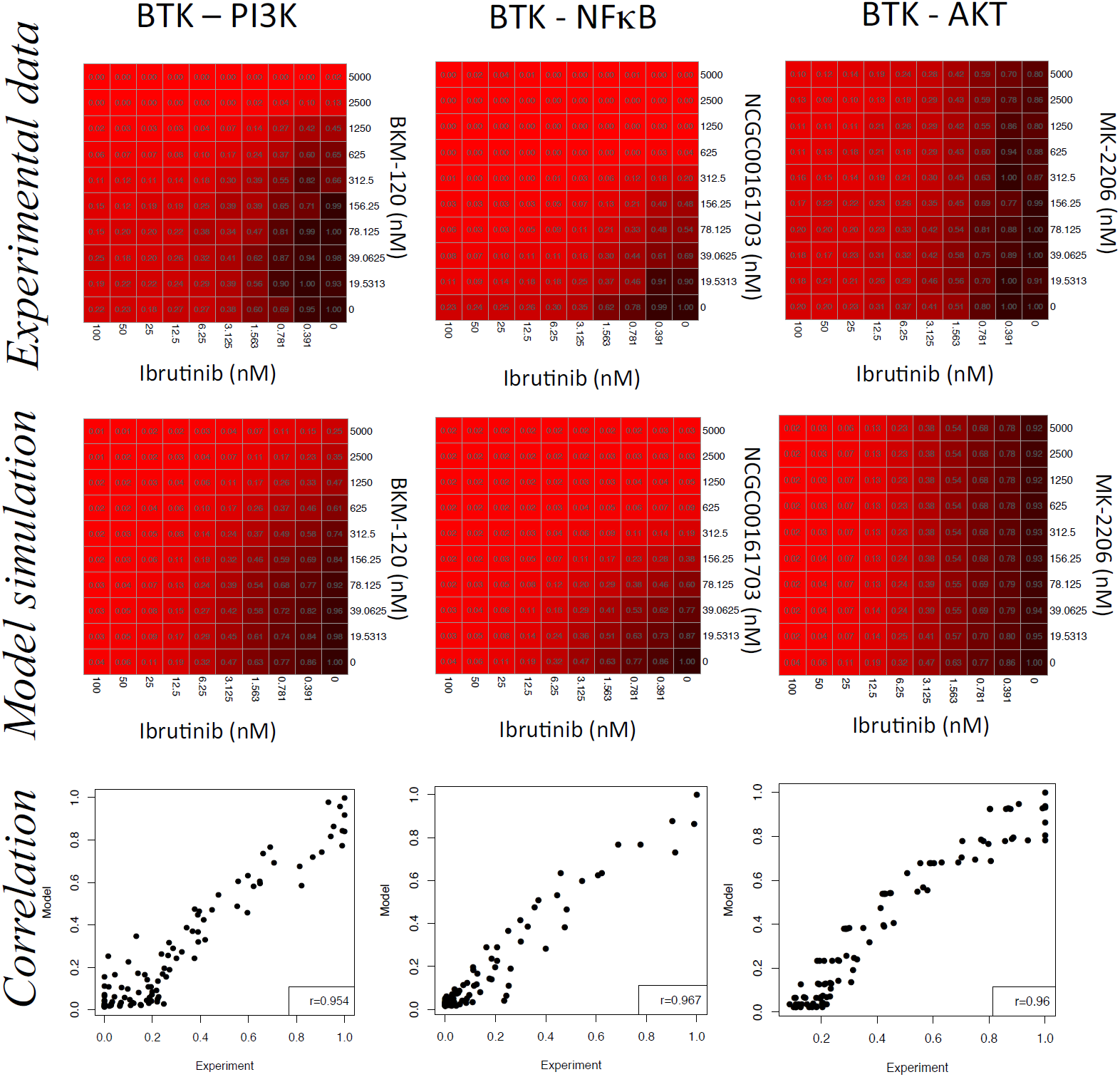
Combinatorial drug viability responses of ibrutinib in combination with additional inhibitors targeting BCR network intermediates were predicted and compared with experimental data.

Overall, these results suggest that viability response of small molecule inhibitors targeting the BCR signaling network can be predicted via *in silico* simulation of the BCR signaling model in combination with the tumor growth model.

### Crosstalk between PI3K and NFκB pathway mediates efficacy of PI3K inhibition in TMD8

In both the drug response data and model’s simulation, we observed that PI3K inhibition is significantly more effective at inhibiting tumor growth than blockage of its downstream effector AKT. A similar phenomenon was reported in other studies, where PI3K inhibition was shown to attenuate NFκB transcriptional activity[3,40]. We hypothesized that the efficacy of PI3K inhibition is primarily attributed to suppression of NFκB signaling, which is mediated by upstream crosstalk between the PI3K and NFκB pathways. To test this hypothesis, we abolished the crosstalk between PI3K and NFκB by knocking out *in silico* PI(3,4,5)P_3_ -mediated membrane recruitment of BTK in the signaling model. Under this condition we re-simulated the viability response of PI3K inhibition, which showed significant reduction compared to both experimental data and simulation with the full signaling model (Fig 4, brown line). This result supports the notion that the upstream crosstalk between PI3K and NFκB pathway is critical in mediating tumor growth inhibition by PI3K inhibitor. It also provides further rational support for the clinical use of PI3K inhibitors in DLBCL that are dependent in NFκB signaling[3,40].

### Computational optimization of targeted therapy against chronic active BCR signaling

Using the above modeling framework, we sought to identify targeted therapies against the BCR signaling network that most effectively inhibit tumor growth. We exhaustively tested all drug pairs based on 11 small molecule inhibitors currently available that target various kinases in the network, yielding 55 treatment strategies in total. In each scenario viability response was simulated at 10 by 10 virtual dosages where each targeted kinase was inhibited at 0% to 99% evenly spaced in log10 space. We calculated area under the combinatorial viability response surface as an overall indicator of drug combination potency. The smaller the value is, the more potent the drug target combination is(Fig 6A). We found that under the same inhibition potency, efficacy of different treatment strategies was highly variable, ranging from almost no growth inhibition to up to 80% reduction (Fig 6B). Specifically, inhibiting downstream of the NFκB signaling pathway, especially through MALT1 and IKK inhibitor, exhibited the most prominent efficacy, and combined MALT1 and IKK blockage yielded highest tumor growth inhibition. In comparison, tumor cell growth was relatively insensitive to blockage of MAPK pathway in our simulations. In summary, this computational screening result suggests that various treatment strategies against a signaling network can yield highly variable therapeutic responses and that in *silico* simulation can help identify targets that confer intrinsic vulnerability.

**Fig 6.**
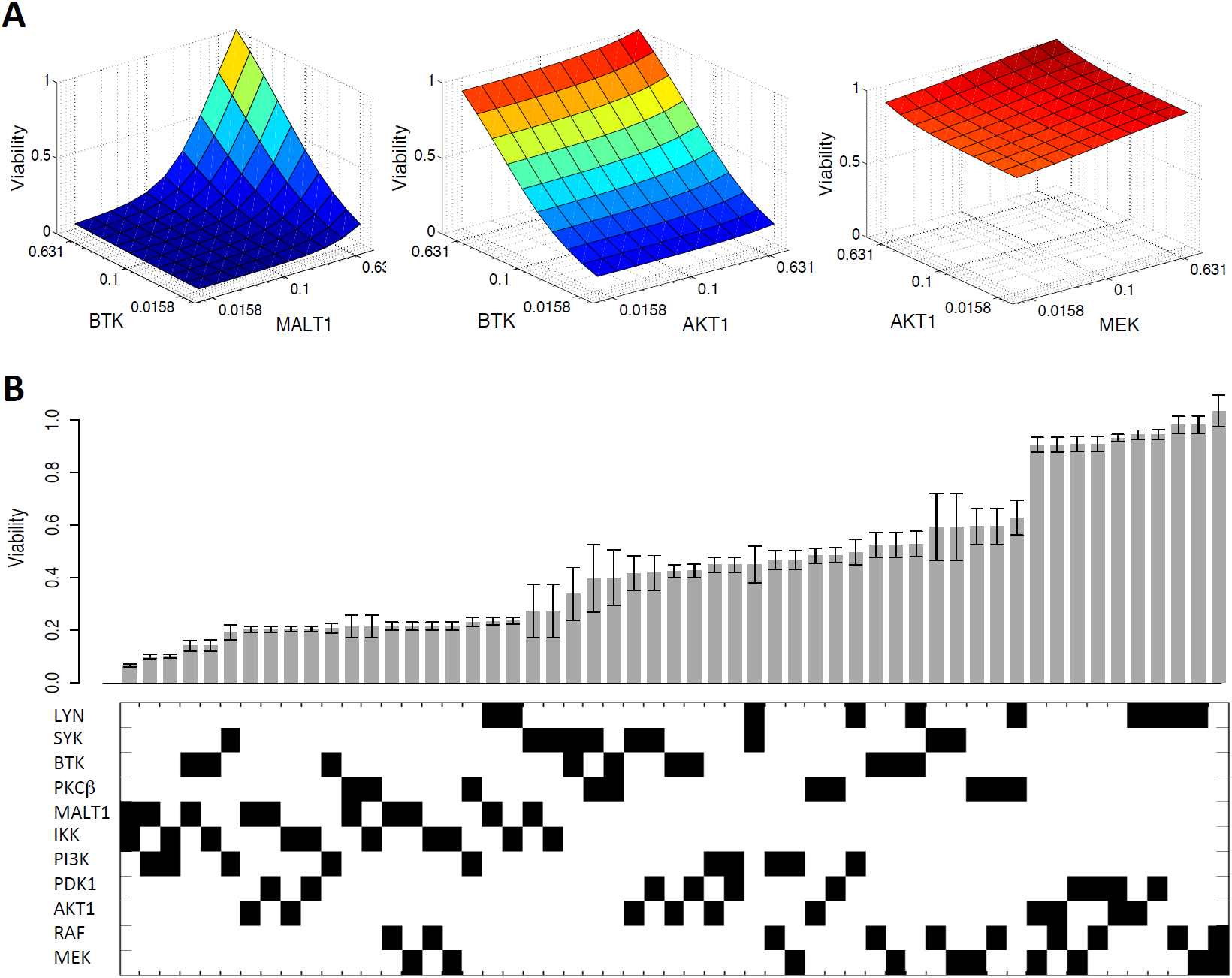
Computational optimization of treatment strategy against chronic active BCR signaling (**A**) Viability response surface of three drug target combination. (**B**) Barplot of simulated viability response of all possible dual inhibition on 11 kinases in the BCR signaling network that are currently targetable. Binary codes on the bottom indicate the treatments applied(black represents targeted inhibition).

We then sought to identify drug combinations that are synergistic via computational simulations. For a given two-drug combination, the combinatorial drug response at 10 by 10 virtual dosage as discussed above were used to estimate mode of drug interaction under the Bliss independence model (**see Materials and Methods**, Fig 7A). Computational screening predicted dual blockage of LYN and SYK as the most synergistic combination. To test this prediction, we treated TMD8 cells with LYN inhibitor Dasatinib and SYK inhibitor R406, at multiple doses. Comparing combinatorial drug response data to theoretical additive response predicted by the Bliss independence model **(see Materials and Methods)**, we confirmed synergism between Dasatinib and R406 (Fig 7B).

**Fig 7.**
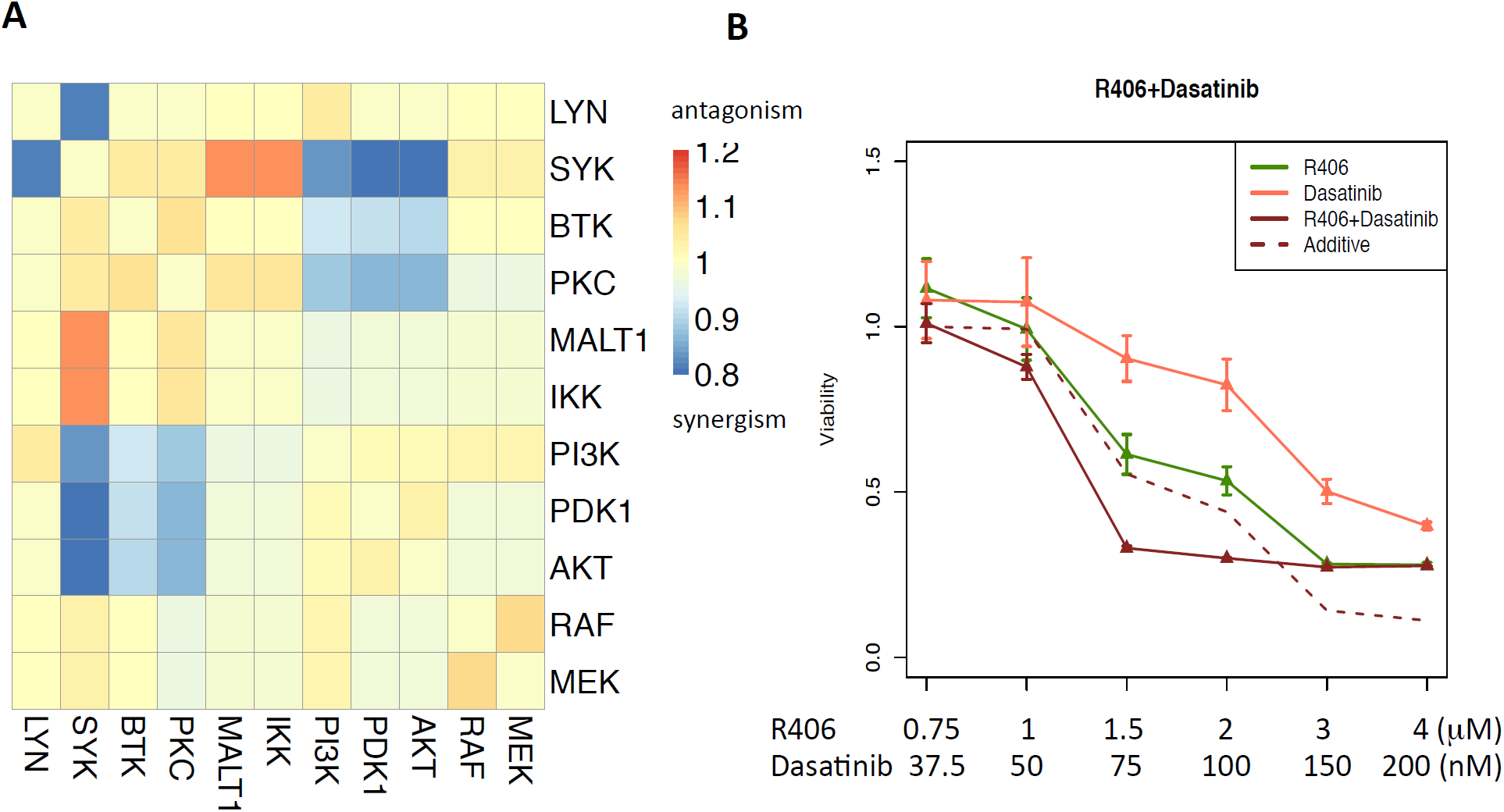
(**A**) Modes of interaction of all pairwise inhibitions under Bliss Independence model. β<1, β=1, β>1 correspond to synergism, additive and antagonism respectively. (**B**) In vitro validation of predicted synergistic drug combination in TMD8. R406 and Dasatinib were inhibitors against SYK and LYN respectively.

## Discussion

It is increasingly acknowledged that aberrant BCR signaling plays a central role in the development and maintenance of many B cell malignancies[41,42]. Though a large panel of small molecule inhibitors against BCR signaling have been developed, rational methodologies that can predict effective combinatorial therapy and guide the design of specific treatment strategy in individual patients have been lacking. We aimed to bridge this gap by constructing the first kinetic model of the core BCR signaling network and using this model to investigate targeted therapy against BCR signaling. We showed that simulations with the signaling model reconstructed dynamics of normal B cell signaling *in silico.* Combining the signaling model with a data-trained tumor growth model successfully predicted viability response of multiple drug combinations, and identified novel synergistic drug combinations such as LYN and SYK inhibitor.

As one of the most important signaling event in B cells, antigen triggered BCR activation has been intensively studied during the past decades. Detailed molecular interactions in the key signal transduction pathway as well as regulatory feedback loops were experimentally identified, providing a unique opportunity to establish a detailed kinetic model of the BCR signaling network. Prompted by the rich information available in literature, we attempted to establish the first kinetic model to quantitatively characterize BCR signaling *in silico.* The model is able to reproduce major kinetic features of BCR signaling observed in experiments. However, we note that simplifications and assumptions in the model may call for further improvements. First of all, how antigen recognition leads to proximal BCR activation, namely phosphorylation of BCR ITAM motif, LYN and SYK, was not addressed in the model. Integration of proximal BCR signaling with downstream signaling model characterized here can potentially provide a more comprehensive understanding of how different strengths of antigen stimulus might lead to various downstream effector activation and distinct cell fate decision. Critically, a negative feedback loop between downstream Ca^2+^ response and upstream phosphatase activity mediated by reactive oxygen species (ROS) may play an important role in determining the threshold and amplitude of BCR response[43]. Furthermore, we did not account for transcriptional regulation of key elements in the BCR signaling network that may influence long term signaling response. Expression of BLNK, CD79A, SYK, BTK, and CD19 is transcriptionally repressed by BLIMP-1, which is activated during germinal center to plasma B cell differentiation triggered by BCR activation[44], Additionally, chemical inhibition of SYK was shown to induce compensatory upregulation of SYK expression mediated by FOXO1[3]. Thus, these transcriptional feedbacks that attempt to upregulate expression of components in the BCR signaling network upon signaling attenuation may mediate resistance to BCR-targeted therapy to some extent.

Oncogenic activation of intracellular signaling pathways drives tumor survival and proliferation by engaging regulators that antagonize apoptosis or drive cell cycle progression. In the BCR signaling network, NFκB transcribes anti-apoptotic factors such as BCL2 and BCR-xL[45] and cell cycle regulators such as cyclin D2[46]. Conversely, AKT and ERK indirectly repress pro-apoptotic factors, e.g., BIM and BAD as well as negative regulators of CDKs such as p27^kip1^ and p21^cip1^[47,48]. A mechanistic characterization of how NFκB, AKT and ERK signal influences tumor survival and proliferation requires deep quantitative knowledge of apoptosis and cell cycle regulation. In this model, we addressed this question by parameterizing a phenotypic tumor growth model from drug response data in TMD8. This parameterization revealed TMD8 to be primarily dependent on NFκB signaling. Under this condition, dual inhibition of IKK and MALT1, two major kinases in the downstream of NFκB signaling cascade, was predicted to have highest growth inhibition efficacy. However, we note that the dependency of various survival and proliferation signals may vary from patient to patient and even dynamically evolve as tumors develop. Indeed, some GCB subtype DLBCL cell lines have been shown to be more sensitive to AKT and ERK inhibition than ABC-DLBCLs [49,50]. When one pathway is blocked by targeted therapy, tumors may adapt by utilizing alternative pathways that remain constitutively active. Consequently, simultaneous repression of all oncogenic pathways, e.g., through dual inhibition of BTK and PI3K, or sequential administration of agents targeting various pathways may ensure more durable response. Monitoring tumor growth and probing signaling dependency for longer periods would help establish mathematical models that can optimize for long-term benefits.

Besides DLBCL, aberrant BCR signaling was shown to play a role in other B cell malignancies such as chronic lymphocytic leukemia(CLL)[51] and mantle cell lymphoma(MCL)[52]. In Phase II studies of BTK inhibitor ibrutinib, 71% and 68% overall response rate(ORR) was reported in CLL and MCL patients respectively[53], suggesting targeting BCR signaling as promising treatment strategy. Correspondingly, predictions reported in this work may be of general guidance for CLL and MCL targeted treatment as well.

## >Materials and Methods

### Cell viability assay

In the published drug response experiment[34], cells were treated at time 0 and incubated for 48 hours. Viability response was normalized to the plate positive control(bortezomib) and negative control(DMSO) as previously described[34]. This normalized data was used for comparison with simulation results.

For our independent validation of synergistic drug combinations, the DLBCL cell line TMD8 was grown in medium containing 90% RPMI and 10% FBS, supplemented with L-glutamine, HEPES and penicillin and streptomycin. R406 and Dasatinib were purchased from Selleck chemicals. Cells were grown at concentrations sufficient to keep untreated cells in exponential growth during the time of drug exposure. Cells were treated with 6 doses of each drug or combination in triplicate. Drug combinations were administered in constant ratio. Cell viability was determined by an ATP luminescent method (CellTiter-Glo, Promega). Luminescence was measured with the Syngery4 microplate reader (BioTek). Cell viability in drug-treated cells was normalized to vehicle treated controls.

### Kinetic model of BCR signaling network

Since protein-protein binding is in general a very fast process at the second time scale, we assumed that this type of reaction were under equilibrium and solved the steady-state level analytically in the model. For a reversible protein-protein binding reaction,

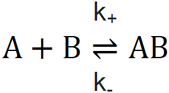

at steady-state,

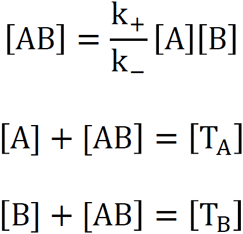

where [A] and [B] stand for the concentration of freed form of A and B; [T_A_] and [T_B_] stand for the total concentration of A and B; [AB] represents the concentration of the complex. By solving the above three equations, we have

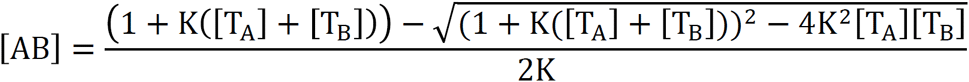

where 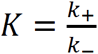, is the inverse of the dissociation constant K_d_.

Under a few circumstances where a protein may bind to more than one partner, the interactions were considered independently for simplification.

For kinase catalyzed reaction, we adopted the classic Michaelis-Menten kinetics,

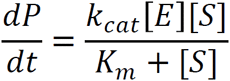

where 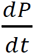 is the rate of catalytic product formation, k_cat_ is the turnover rate, K_m_ is the Michaelis-Menten constant, [E] and [S] are concentration of enzyme and substrate respectively.

Reactions in the BCR signaling network were written into corresponding equations according to rules discussed above. The full model consists of 28 state variables each representing concentration of a specific form of a protein species, depicted by 10 steady-state equations and 18 ODEs (**see S1 Text**). Parameters of total protein concentrations are summarized in **S3 Table**. Kinetic parameters in the model are summarized in **S4 Table**. We performed parameter sensitivity analysis where each parameter was perturbed independently across four orders of magnitudes and viability response was recorded. We found overall robustness and identified the most sensitive parameters as parameters regulating main axis of the NFκB pathway (**see S1 Fig**).

### Input signal

We imposed temporal pLYN and pSYK stimulus in normal BCR signaling modeled by two double exponential functions,

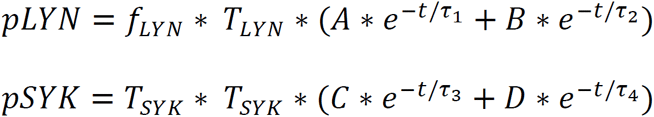

where T_*LYN*_ and T_*SYK*_ are total concentration of LYN and SYK respectively. f_*LYN*_ and f_*SYK*_ are the phosphorylated fraction of LYN and SYK respectively. f_*LYN*_ and f_*SYK*_ were estimated by fitting to the phosphorylation time course data using genetic algorithm(Fig 3). A,B,C,D τ_1_, τ_2_, τ_3_, τ_4_. were estimated by fitting to the normalized pLYN and pSYK time course.

In contrast, we imposed constitutive pLYN and pSYK stimulus in simulations of diseased ABC-DLBCLs, with negative feedback from SYK and PKC respectively.

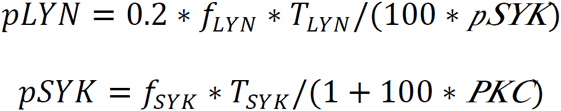

*The* 0.2 coefficient is to account for LYN attenuation effect due to CD79B mutation in TMD8[2]. The parameter for negative feedback is estimated by fitting to single drug viability response.

### Tumor growth model

Assume a tumor cell population is at exponential growth phase,

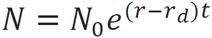

where *r*_d_ is basal death rate, while growth rate *r* is dependent on three downstream survival and proliferation signals NFκB, pAKT, and pERK (normalized by untreated control),

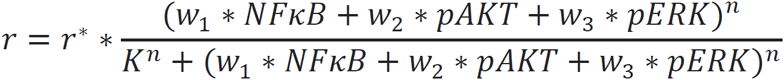

Therefore, viability response defined as the ratio of cell number monitored under treated condition *N* (for a time span of *T*) and untreated control *N_c_* can be predicted as following,

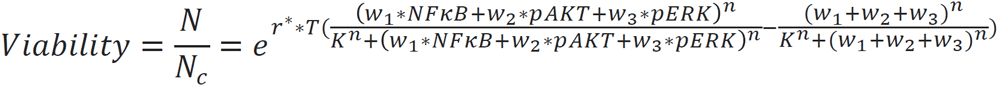

Parameters required in this function were trained with viability data of three single drug viability responses, namely NFκB, AKT and MEK inhibitor respectively. First the level of the three downstream survival and proliferation signals were predicted via simulation of the signaling model, and then input into the tumor growth functions to compute the viability output. Parameters were chosen by minimizing the sum of residuals between the viability prediction and experimental data.

### Drug kinetics

To simulate an inhibitor’s effect at a given dosage, percent activity of the targeted kinase was calculated via the medium effect equation,

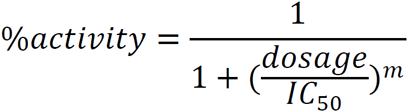

The drug’s IC50 was taken from literature (**S2 Table**), while m was assumed to be 1 under a first order approximation. Then the activity of the targeted kinase (i.e. parameters representing catalytic or activation rate of targeted kinase) was reduced to the corresponding percentage in the kinetic model. We list perturbed parameters in each simulated inhibitor treatment in **S2 Table**.

### Synergy quantification

Under the Bliss Independence model, the additive effect of two inhibitors is computed as the multiplication of the effect of individual inhibitors,

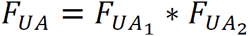

where F_*UA*_ indicates the fraction unaffected. To evaluate mode of interaction between two inhibitors, we computed viability response at 10x10 virtual dosages by varying the percent inhibition of each targeted kinase independently from 0% to 90% at 10% interval. These viability values were used to estimate parameter that minimizes the following metric,

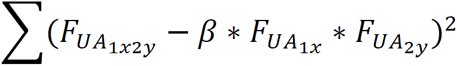

where x,y are virtual dosages for inhibitor 1 and 2 respectively. *β*< 1, *β* = 1, *β* > 1 indicates synergism, additive and antagonism respectively.

## Acknowledgement

This work was supported by NIH, CAREER grant from National Science Foundation, Starr Cancer Consortium, and Hirschl Trust. AM is supported by NCI R01 CA143032, R01 CA104348, the Burroughs Welcome Foundation, and the Chemotherapy Foundation. We thank all Elemento and Melnick lab members for productive discussions.

## Supporting Information

S1 Text. Kinetic model equations.

S1 Table. Parameter bounds.

S2 Table. Inhibitor against BCR signaling network.

S3 Table. Total protein concentration.

S1 Figure. Parameter sensitivity analysis.

S4 Table. Kinetic model parameters.

